# Identification of Biomarkers and Trajectories of Prostate Cancer Progression: A Bioinformatics Fusion of Weighted Correlation Network Analysis and Machine Learning

**DOI:** 10.1101/2023.03.02.530740

**Authors:** Raheleh Sheibani-Tezerji, Carlos Uziel Pérez Malla, Gabriel Wasinger, Katarina Misura, Astrid Haase, Anna Malzer, Jessica Kalla, Loan Tran, Gerda Egger

## Abstract

**Background:** Prostate cancer diagnosis and prognosis is currently limited by the availability of sensitive and specific biomarkers. There is an urgent need to develop molecular biomarkers that allow for the distinction of indolent from aggressive disease, the sensitive detection of heterogeneous tumors, or the evaluation of micro-metastases. The availability of multi-omics datasets in publicly accessible databases provides a valuable foundation to develop computational workflows for the identification of suitable biomarkers for clinical management of cancer patients.

**Results:** We combined transcriptomic data of primary localized and advanced prostate cancer from two cancer databases. Transcriptomic analysis of metastatic tumors unveiled a distinct overexpression pattern of genes encoding cell surface proteins intricately associated with cell-matrix components and chemokine signaling pathways. Utilizing an integrated approach combining machine learning and weighted gene correlation network modules, we identified the EZH2-TROAP axis as the main trajectory from initial tumor development to lethal metastatic disease. In addition, we identified and independently validated 58 promising biomarkers that were specifically upregulated in primary localized or metastatic disease. Among those biomarkers, 22 were highly significant for predicting biochemical recurrence. Notably, we confirmed TPX2 upregulation at the protein level in an independent cohort of primary prostate cancer and matched lymph node metastases.

**Conclusions:** This study demonstrates the effectiveness of using advanced bioinformatics approaches to identify the biological factors that drive prostate cancer progression. Furthermore, the targets identified show promise as prognostic biomarkers in clinical settings. Thus, integrative bioinformatics methods provide both deeper understanding of disease dynamics and open the doors for future personalized interventions.

## 1. Background

Prostate cancer (PCa) is the second most common cancer in men with 1.41 million cases worldwide in 2020, accounting for an incidence of 23.6 and mortality of 3.4 per 100,000 cases [1]. Risk factors include age, family history, race, germ line mutations and clinical predisposition (e.g. Lynch syndrome) [1]. Clinically, PCa ranges from indolent and slowly progressive disease, which can be cured with surgery and/or radiation therapy, to aggressive disease including metastatic castration-resistant PCa (mCRPC) [2]. Approximately 20-40% of patients experience biochemical recurrence (BCR) defined by rising prostate specific antigen (PSA) levels after radical prostatectomy due to local recurrence or metastasis. Metastasis most frequently involves bone, distant lymph nodes, liver and lungs and represents the main cause of disease related mortality.

Currently, only a handful of PCa biomarkers are in clinical use. The most prominent is PSA, which is used for screening, diagnosis, monitoring and risk prediction of PCa. PSA screening has limitations due to overdiagnosis, leading to unnecessary prostate biopsies and aggressive treatment regardless of the risk [3]. Additionally, prostate-specific membrane antigen (PSMA), a transmembrane protein, is highly overexpressed in PCa (100-to 1000-fold) and is a theranostic target of PCa, both for diagnosis and treatment in nuclear medicine [4]. Expression levels of PSMA are positively correlated with more aggressive disease, high PSA, high Gleason scores and early recurrence.

PCa risk stratification at diagnosis and treatment decisions are currently based on clinical parameters including Gleason score, PSA level and tumor staging. In addition, several risk stratification methods to predict BCR, metastasis or for therapy stratification have been tested and validated based on gene expression signatures, some of which are already in clinical use [5–7].

Numerous studies have performed meta-analyses of publicly available datasets and databases such as The Cancer Genome Atlas (TCGA) [8]. Aside from biomarker discovery, access to public data provides significant insight into understanding biological pathways, modifications and functions of established biomarkers in certain patients. *In silico* data mining is a useful tool to decipher relevant disease biomarkers and to generate research hypotheses. In particular, various machine (deep) learning algorithms have been developed and also applied to PCa data, which have high potential to interrogate large amounts of multi-dimensional data [9–13].

Here, we integrated publicly available gene expression datasets of localized PCa with matched adjacent normal tissue from The Cancer Genome Atlas (TCGA) prostate adenocarcinoma (PRAD) dataset [8] with mCRPC samples from two recent studies [14, 15], which were accessed via the database of Genotypes and Phenotypes (dbGAP) [16]. In order to identify prognostic biomarkers for PCa development and disease progression to metastasis, we performed pair-wise differential gene expression analysis identifying deregulated genes between localized primary PCa and normal adjacent tissues, as well as between primary PCa and mCRPC specimens. Differentially expressed genes were subsequently used to infer significant targets based on several computational methods including pathway analysis, random forest machine learning, weighted correlation network analysis (WGCNA) and logistic regression models. A distinct focus was set on genes that encode for cell surface and extracellular proteins, in order to identify potential diagnostic and therapeutic targets. Using this strategy, we identified several well established prognostic PCa biomarkers in addition to novel targets, which showed association with DNA damage repair pathways or cell cycle and were correlated to disease-free survival of patients. These markers might represent important biological drivers of PCa progression and prove useful as diagnostic and prognostic markers as well as therapeutic targets in the future.

## 2. Methods

### RNA-Seq data preparation

Data used in this study originated from two data cohorts extracted from the cancer genome atlas (TCGA) [8] and the database of Genotypes and Phenotypes (dbGAP) [16]. The first set of RNA-Seq data belongs to the TCGA Prostate Adenocarcinoma project (TCGA-PRAD, https://portal.gdc.cancer.gov/repository, Last accessed in 09/2022). We downloaded the TCGA raw STAR read counts of prostate cancer tumor samples from the GDC data portal using the R/Bioconductor package TCGAbiolinks [17]. The PRAD dataset includes the entire collection of 500 primary solid tumors and 52 solid tissue normal samples.

For the second set of RNA-Seq data, we obtained authorized access to the metastatic castration resistant prostate cancer (mCRPC) tissue samples of the SU2C-PCF dataset [14, 15] via dbGAP, with accession number phs000915.v2.p2 (https://www.ncbi.nlm.nih.gov/projects/gap/cgi-bin/study.cgi?study_id=phs000915.v2.p2).

SU2C-PCF comprises two studies, where a total of 429 patients were enrolled at an international consortium, all of whom underwent biopsy for the collection of mCRPC tissue. RNA sequencing was performed on a subset of 224 tumors.

We obtained and processed the .sra files of all SU2C-PCF samples, as well as the sample annotation information coming from different sources, such as basic annotation from dbGAP, clinical annotation from cBioPortal [18] (https://cbioportal-datahub.s3.amazonaws.com/prad_su2c_2019.tar.gz) and SRA metadata from NCBI (https://0-www-ncbi-nlm-nih-gov.brum.beds.ac.uk/Traces/study/?acc=phs000915&o=acc_s%3Aa).

Data exploratory analysis clearly showed that metastatic samples were perfectly clustered into two groups after applying K-means to the PCa results. An in-depth analysis of the available data annotation showed that samples were separated based on the annotation for “sample use” as either “RNA_Seq” or “RNA_Seq_Exome”. For this reason, and due to their closer similarity to the TCGA samples, we only included samples labeled as “RNA_Seq” in our study. Additionally, we removed all sample replicates per group, keeping only those samples that were sequenced the latest (samples sequenced in 2018 and 2019 at the University of Michigan, Broad Institute). A total of 14 samples with neuroendocrine features were also excluded from our study. Therefore, of the 224 RNA-Seq samples available on dbGAP [14, 15], we finally included 57 samples with unique patient IDs to avoid technical bias (Metastatic_BB in additional file 01). These samples were obtained from biopsies of lymph node (n=28; 50.90%), bone (n=13; 23.63%), liver (n=8; 13.54%), and other soft tissues (n=6; 7.27%).

### Pre-processing

While the raw counts were directly available for normal and primary tumor samples from TCGA, mCRPC samples from dbGAP required preprocessing. We extracted the fastq paired-end reads from the sra files and used cutadapt [19] to remove unwanted sequences (e.g., adapters, poly-A tails, etc.) and low-quality reads. Then, the STAR aligner tool [20] was used to map the reads to the GRCh38.p13 (release 105) human reference genome, obtained from ENSEMBL (https://www.ensembl.org/Homo_sapiens/Info/Index), as well as to obtain gene counts.

Only intersecting sequenced transcripts of TCGA and dbGAP count files were kept, resulting in 60,528 common transcripts. We also performed a naive pre-filtering step to remove low count genes by only keeping those transcripts with an average count across all samples bigger than 1. This yielded a total of 33,010 transcripts for further analysis.

### Transcriptome analysis

We used the R/Bioconductor package DESeq2 [21] to perform differential gene expression (DE) analysis of the un-stranded STAR transcript counts. Differentially expressed genes (DEGs) were derived from likelihood ratio tests in the respective experimental subgroups: primary versus normal (prim/norm) and metastatic versus primary (met/prim). Genes with an adjusted P-value (Padj) < 0.05 and absolute log2 fold change (LFC) > 1 were considered significantly differentially expressed, representing a conservative and stringent approach. Finally, read counts were normalized by the DESeq2 normalization method of variance stabilizing transformation (VST), to be used in downstream analyses.

### Functional gene set enrichment analysis for differentially expressed genes

To gain insight into the biological and/or clinical relevance of the DEGs in each experimental subgroup, enrichment analysis was performed using several tools, including GO enrichment analysis, cell surface protein identification, biomarker prediction and pathway analysis.

The expression patterns of DEGs in each pair-wise comparison (prim/norm and met/prim) were explored by hierarchical clustering (hclust method) using the R package simplifyEnrichment [22]. The classification of samples was obtained by consensus partitioning with the R package cola [23]. The signature genes were additionally clustered into two groups by k-means clustering (cluster 1, cluster 2). Furthermore, GO enrichment analysis was applied to the two groups of genes separately with the R package clusterProfiler [24]. To visualize whether the GO terms were significantly enriched in each subgroup, the binary cut was applied directly to the union of the two significant GO term lists, and a heatmap of Padj was placed on the left side of the GO similarity heatmap (Padj < 0.05). This strategy keeps all significant GO terms without removing any. The summaries of the biological functions in clusters are visualized as word clouds and are attached to the GO similarity heatmap, which gives a direct illustration of the common biological functions involved in each cluster.

### Pathway analysis

With the help of the R package clusterProfiler [24], we mined enriched pathways from Reactome databases [25]. Additionally, we employed ingenuity pathway analysis (QIAGEN IPA (QIAGEN Inc., https://digitalinsights.qiagen.com/IPA) [26] to interpret the DEGs in the context of biological processes, pathways and networks using the ’’core analysis” function. The “biomarker analysis” function included in IPA® was used to identify targets, which were already reported as biomarkers in previous studies.

### Cell surface protein analysis

We performed cell surface protein (CSP) analysis using the cell surface protein atlas (CSPA), a public resource containing experimental evidence for cell-surface proteins identified in 41 human cell types [27]. The basis and reference for the presented human surface proteome analysis was the human proteome in UniProtKB/Swiss-Prot (Version 2015_01) [28]. We matched the differentially expressed genes to the human peptides from the CSPA, generating a list of cell surface proteins enriched in our data.

To gain insight into the functional aspects of up-regulated cell surface receptors (n=201) in the met/prim comparison, the respective genes were used as input for integrative pathway analysis [29]. Pathways and gene sets from the Gene Ontology (GO), Human phenotype ontology (HP), WikiPathways (WP) and the Reactome collections were downloaded from the g:Profiler web server [30] as a GMT file.

Active pathways were used as input for the EnrichmentMap app [31] of Cytoscape [32] for network visualization of similar pathways. Then, we used the AutoAnnotate app [33] to summarize the networks and clustering pathways. Auto-annotated enrichment maps of up-regulated CSP receptors were visualized with stringent pathway similarity scores (Jaccard and overlap combined coefficient 0.6) and manually curated for the most representative groups of similar pathways and processes. Pathways that were redundant with larger groups of pathways were merged with the latter or discarded. The coloring of pathways was done according to their g:Profiler supporting resources.

### DEGs filtering with machine learning

Explainable machine learning (ML) models can be a useful tool to rank DEGs based on their ability to predict different conditions. We trained ML models on our DEGs’ normalized expression data and used them to produce “importance scores” for each DEG.

Input gene expression data was pre-processed in multiple steps. First, we filtered our DEGs of interest (Padj < 0.05, LFC > 1) by keeping only those transcripts that were uniquely mapped to ENTREZID/SYMBOL and were either protein-coding or non-coding RNA. We also removed transcripts that mapped to the same ENTREZID. For met/prim, we also removed genes whose value ranges did not overlap between classes. This resulted in 1,406 input DEGs for prim/norm and 3,788 for met/prim. As a last step, we built the input matrix using the variance-stabilized transformed (VST) expression counts of our filtered DEGs and performed min-max feature scaling.

We trained our models following two main steps. First, we obtained the optimal hyper-parameters by applying grid search cross-validation on our data, as implemented in the Python package scikit-learn [34]. Second, we performed bootstrap training for 10.000 iterations. In other words, we trained each model multiple times, each under a different random seed, which impacts the stochasticity of the training, from model fitting to train/test (80/20) data splits. For each bootstrap training iteration, we calculated the shapely additive explanations (SHAP) [35] values for all samples, a method based on cooperative game theory used for interpretability of ML models. SHAP values were then averaged among all iterations, resulting in the final gene importance scores for a given contrast.

Finally, we used the DEGs SHAP values in each experimental subgroup to rank the genes as well as to keep only the most relevant ones (i.e., those with SHAP values > 0.001).

### WGCNA

In order to obtain additional insights from our DEGs, we explored the co-expression relationships among them with the R/Bioconductor package WGCNA [36].

We first loaded the VST expression counts and pre-processed our data the same way we did for the ML analysis, except that we did not remove non-overlapping genes and avoided any post-processing scaling. This resulted on 1,406 input DEGs for prim/norm and 3,804 for met/prim.

We then detected modules of genes as well as their hub genes, based on node degree within each module. For this, we first did a power estimate to weight the co-expression network to approximate a scale free topology, using a signed network type and biweight midcorrelation as correlation type. Furthermore, we mined all the modules detected in these analysis via functional enrichment pathways for Reactome databases [25] using the R package clusterProfiler [24].

### Final evaluation of potential biomarkers

We obtained sets of potential biomarkers by intersecting the up-regulated DEGs obtained from the different methods (i.e. DE, ML and WGCNA) for both of our experimental subgroups. We performed an in-silico validation of these biomarkers on an external dataset, the Prostate Cancer Transcriptome Atlas (PCTA) [37]. This dataset contains 174 normal, 714 primary and 316 metastatic samples from 11 datasets. We removed all overlapping samples from TCGA-PRAD and SU2C-PCF and kept only the common transcripts. This step yielded a dataset with 122 normal, 209 primary and 180 metastatic samples. We ran our DE pipeline on this data to obtain the log2 fold change and P-value adjusted metrics. Then we examined whether our biomarkers behaved similarly for each experimental subgroup when comparing PCTA to our data. That is, whether we could observe the same up-regulation patterns of our pre-selected biomarkers. Lastly, we performed recursive partitioning-based survival analysis [38] of the 58 individual biomarkers using the MSKCC dataset [39] via the web-based camcAPP tool [40].

### Generation of tissue micro arrays and immunohistochemistry analysis

PCa tissues for validating immunohistochemical (IHC) staining were collected at the Department of Pathology (Medical University of Vienna). Formalin-fixed paraffin-embedded (FFPE) samples of 51 primary PCa (of which 25 were multifocal) and 35 matched lymph node metastases were utilized to generate 3 tissue microarrays (TMA) containing a total of 402 cores, including 194 primary PCa cores, 154 benign prostate gland cores, and 54 metastasis cores. TMA FFPE blocks were cut into 3 µm thick sections and manual IHC staining was performed using a Rabbit Anti-Human TPX2 antibody (11741-1-AP, Proteintech) diluted at 1:200. Counterstain was performed using hematoxylin. Interpretation of marker expression in tissue samples was performed by a pathologist trained in uropathology, using high-resolution brightfield scans performed using the Vectra Polaris™ Automated Quantitative Pathology Imaging System by Akoya Biosciences® at 40x magnification. H-scores were calculated for each primary tumor focus, metastasis and benign prostate tissue by estimating the average % of nuclear expression in all cores of one compartment and multiplying this value by a factor based on the staining intensity (x1 for weak staining, x2 for moderate staining, x3 for strong staining), resulting in a H-score ranging from 0-300. Estimation of an average H-score between all available cores in the final IHC slides resulted in analysis of a total of 75 primary PCa foci, 51 matched benign prostate tissues, and 32 matched metastatic tissues.

The source code used for our analysis are available in a GitHub repository for reproducibility and further research (https://github.com/CarlosUziel/pca_wgcna_ml).

## 3. Results

To infer PCa specific gene expression signatures that reflected general transcriptional trajectories of PCa development and progression irrespective of molecular subtype (see analysis workflow in Figure 1A), we considered RNA-Seq data of 500 primary prostate adenocarcinomas with 52 matched adjacent normal tissue samples from the PRAD dataset of the TCGA [8], and 57 mCRPC samples from the dbGAP databases, which were recently published [14, 15] (Figure 1B and Additional file 02). Metastatic samples were derived from different sites including lymph nodes, bone, liver and other soft tissues (Figure 1C).

**Figure 1.**
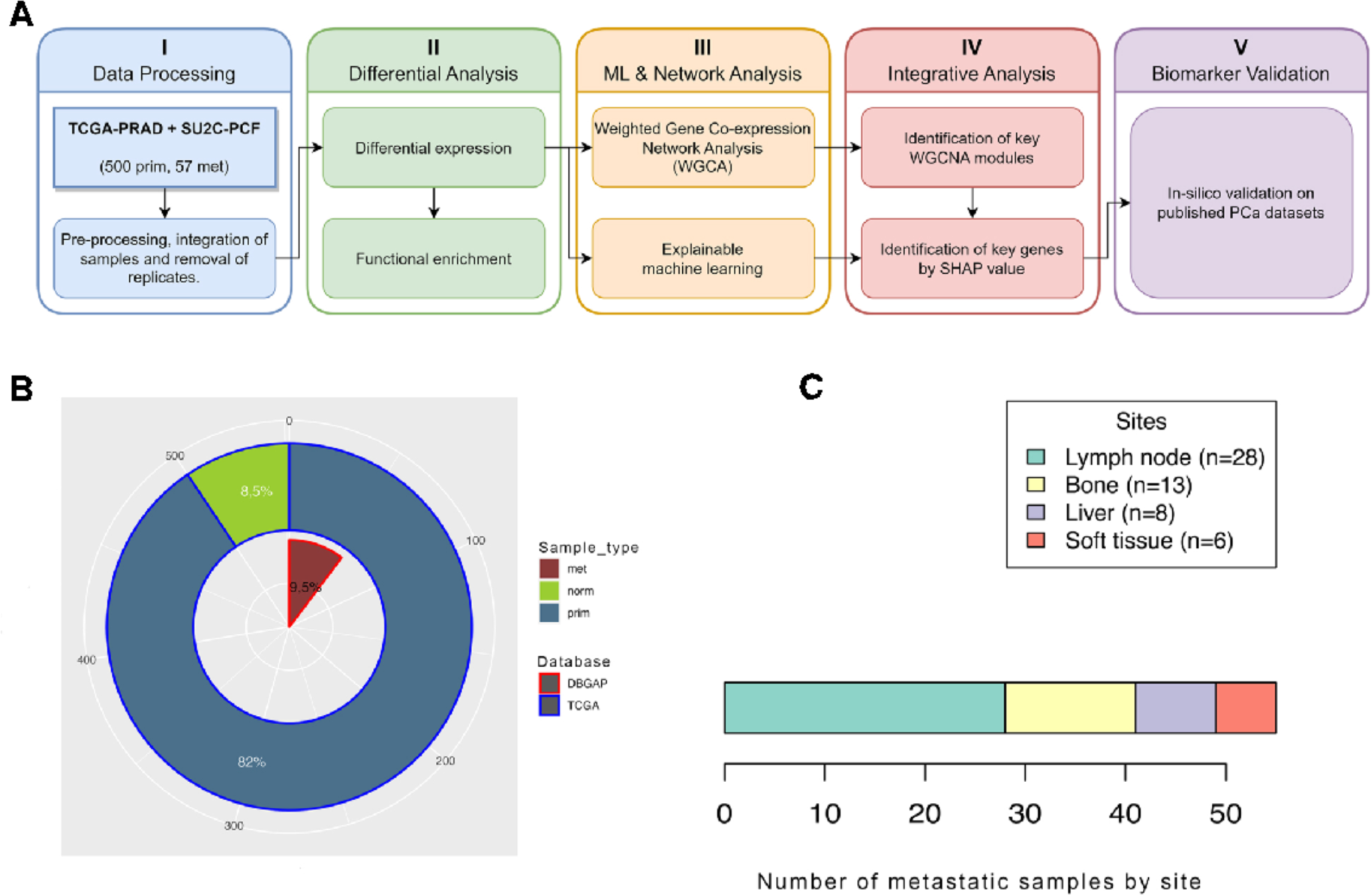
Analysis setup. **(A)** Schematic presentation of the analysis workflow. This diagram shows the 5 principal steps of analysis procedure, which are data processing, differential analysis, ML and network analysis, integrative analysis and biomarker validation. **(B)** Overview of sample origin and grouping for metastatic (n=57), primary tumor (n=500) and normal adjacent (n=52) samples from patients with PCa and mCRPC obtained from TCGA [8] and dbGAP[14, 15] databases. **(C)** Number and biopsy sites of mCRPC samples. (met, metastatic; norm, normal adjacent tissue; prim, primary).

### Global scaling of gene expression profiles unveils (dis)similarities between normal and tumor samples

In order to assess the relations of the different sample types based on their gene expression signatures, we applied principal component analysis (PCA), which confirmed that most of the variation in expression levels was due to the different sample groups. We observed a clear separation of metastatic (met) samples from primary (prim) and normal adjacent (norm) samples, with some degree of overlap noted between norm and prim tissues, accounting for approximately 22% of the data variance of the two major principal components (PC1=13%, PC2=9%) (Figure 2A). The remaining components were sorted in descending order of their contribution to the variance, (PC3=7%, PC4=4%) (Additional file 03). Additionally, unsupervised hierarchical clustering of normalized gene expression data also showed a clear separation of normal and metastatic samples from primary tumors (Figure 2B).

**Figure 2.**
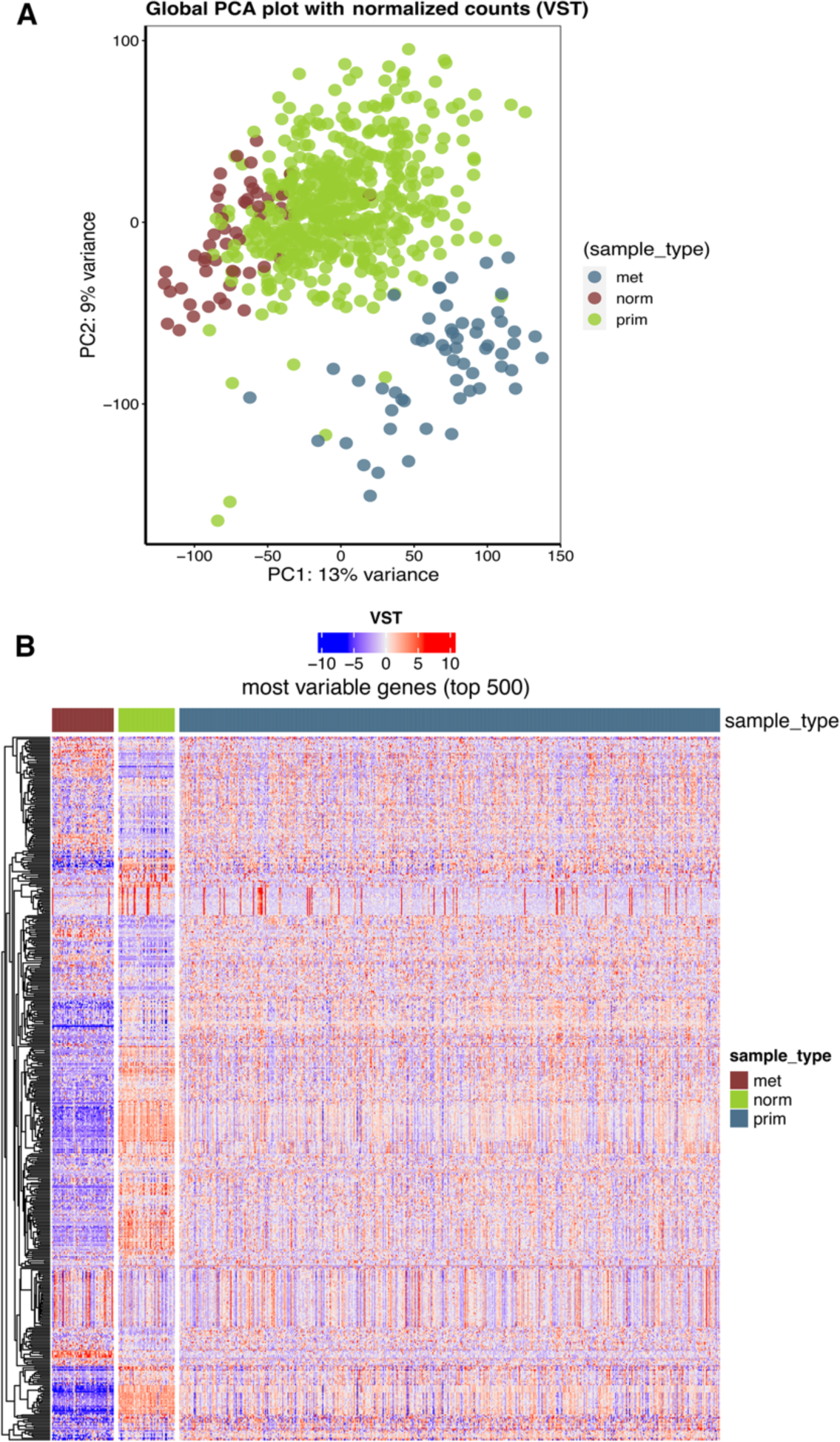
Global scaling of RNA-Seq data. **(A)** Principal component analysis (PCA) of RNA expression levels. The first two components shown explain the largest part of the variation in RNA expression (22%). Individual samples (circles) are colour-coded by sample type (met, metastatic; norm, normal adjacent; prim, primary). **(B)** Heatmap of the unsupervised hierarchical clustering analysis on the variance-stabilized transformed (VST) expression of the top 500 most variable genes across all samples within each sample type. Each row corresponds to a single gene, whereas each column corresponds to a single sample.

### Identification of differentially expressed genes

To gain a comprehensive insight into the transcriptional changes among the norm, prim, and met sample groups, we performed pairwise differential gene expression (DE) analysis on the combined TCGA and dbGAP datasets. For consecutive downstream analyses, we focused exclusively on differentially expressed genes (DEGs) with unique gene IDs (e.g., ENTREZID) and, unless otherwise specified, filtered by Padj < 0.05 and absolute LFC > 1.

The analysis of prim/norm samples resulted in 3,554 DEGs, 1,670 of which were up and 1,884 were down-regulated (Supplementary Table S1, Additional file 04). The met/prim comparison identified 6,230 significantly deregulated genes, including 4,532 up and 1,698 down-regulated genes (Supplementary Table S2, Additional file 04). Notably, we observed a subset of 606 up and 350 down-regulated DEGs that were shared between prim/norm and met/prim, respectively, suggesting their potential importance in both tumor initiation and progression (Figure 3A, Supplementary Tables S3-S4, Additional file 04).

**Figure 3.**
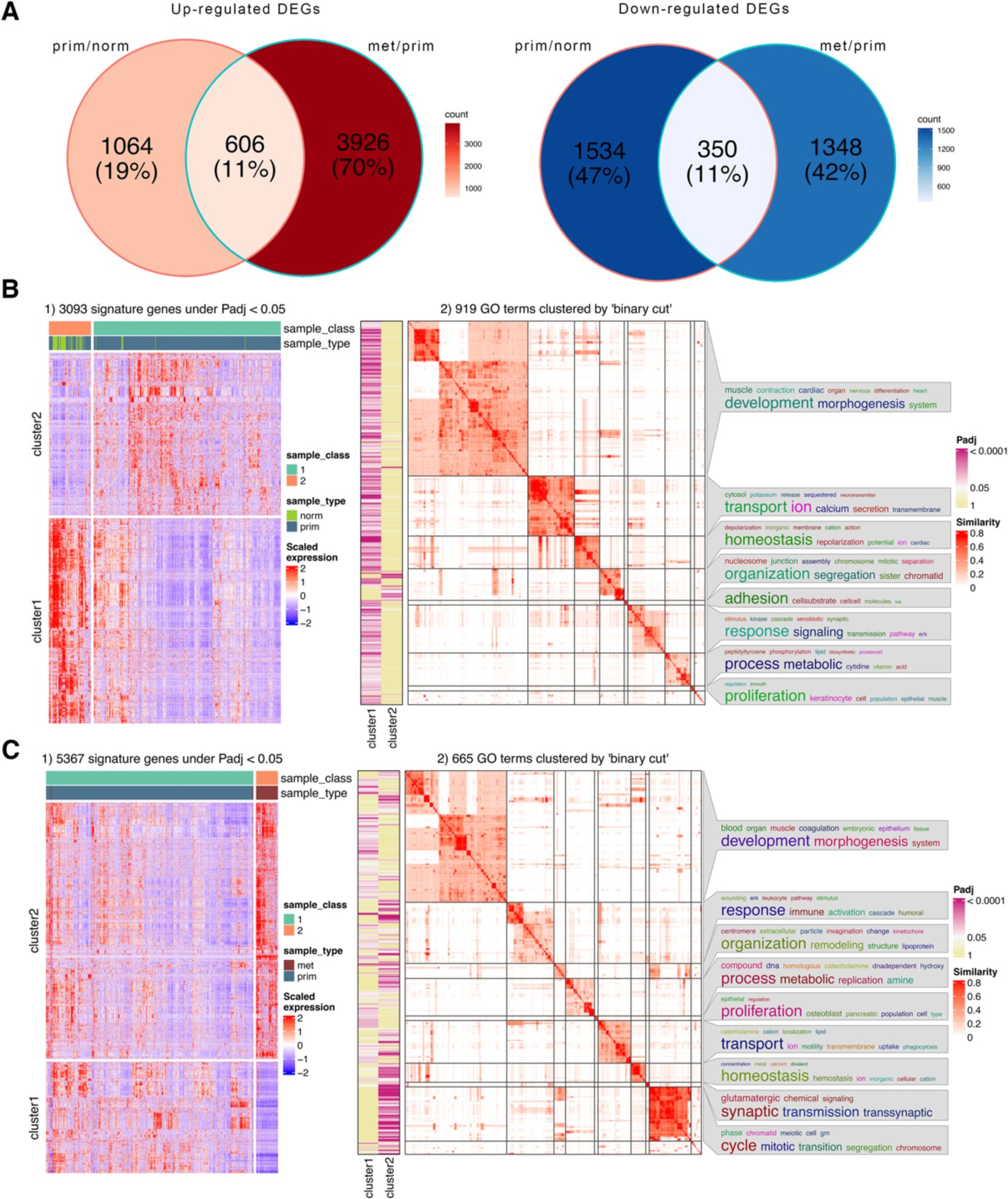
Expression patterns and biological function of differentially expressed genes. **(A)** Venn diagrams illustrate overlaps and unique features identified in up and down-regulated differentially expressed genes (DEGs) in each pair-wise comparison (prim/norm; met/prim). and **(C)** Hierarchical clustering of signature genes differentially expressed between prim/norm and met/prim samples (B1 and C1). Rows correspond to genes and columns correspond to samples. Sample class was obtained by consensus partitioning methods to separate samples into subclasses. The two gene clusters (cluster1 and cluster2) were generated by applying k-means clustering on rows of the expression matrix. The z-score standardization was applied on matrix rows. **(B)** and **(C)** Significant GO terms enriched in either of the corresponding gene clusters were grouped based on their similarities and visualized as a heatmap (B2 and C2). The columns on the left of the GO heatmap indicate the significance levels of GO terms in the individual clusters (Padj < 0.05). The summaries of the biological functions in clusters are visualized as word clouds and are attached to the GO similarity heatmap. In word clouds, enrichment of keywords is assessed by Fisher’s exact test, and the significance is mapped to the font size of keywords.

To define the top gene expression signatures specific for each sample type, we performed hierarchical clustering analysis using DEGs identified from pairwise comparisons of prim/norm and met/prim samples. We employed consensus partitioning to group samples into two distinct classes. For the prim/norm comparison, some normal samples overlapped with primary tumors in one class, whereas the metastatic samples were more clearly separated from the primary tumors in individual classes. The DEGs with zero or low variance variance (sd <= 0.05 quantile) were filtered out, resulting in 3,093 and 5,367 signature genes from prim/norm and met/prim, respectively. These signature genes were then grouped into two separate gene expression clusters in both analyses (Figure 3B1-C1).

To explore the systematic features and biological functions of the differentially expressed signature genes, gene ontology (GO) enrichment analysis was applied to the two gene clusters of prim/norm and met/prim (Additional file 05). Significant GO terms (Padj < 0.05) were then clustered based on their similarities. Furthermore, GO enrichment results were visualized as word clouds to summarize the biological functions in each cluster (Figure 3, B2 and C2). Down-regulated genes from both prim/norm and met/norm were associated with developmental pathways, whereas up-regulated genes from both pairwise comparisons were associated with GO terms related to cell cycle and mitosis. Furthermore, metabolic processes and signaling pathways were enriched in up and down-regulated gene expression clusters of both comparisons. Together, these data indicate that similar pathways are affected by differentially expressed genes in the transition from normal to primary tumors and to metastatic disease.

### Functional gene set association analysis of DEGs

To identify candidate diagnostic or prognostic PCa biomarkers as well as putative therapeutic targets, we subsequently focused our analyses only on up-regulated DEGs (Padj < 0.05 and LFC > 1) from the pairwise comparisons prim/norm and met/prim. Ingenuity Pathway Analysis (IPA) [26] identified several known PCa signaling pathways including TGF-ß, p38-MAPK, cAMP and PKA signaling in prim/norm samples and cAMP, PKA, PI3K-AKT, CREB and G-protein coupled receptor signaling in the met/prim analyses (Additional file 06). Using the biomarker analysis tool included within the IPA software, a total of 34 genes were identified from the up-regulated DEGs of prim/norm samples and 114 from the up-regulated DEGs of met/prim tumors, 21 of which were found in both comparisons (Additional file 07). Among the biomarker candidates we found cytokines and growth factors, enzymes and peptidases, transmembrane receptors and transporters as well as transcription regulators and other mostly secreted factors. Interestingly, several factors associated with innate immunity and acute phase response proteins including albumin, C-reactive protein, fibrinogen, prothrombin, or haptoglobin were up-regulated in primary and metastatic PCa. Some of these factors were already suggested as biomarkers for non-invasive diagnostics in liquid biopsies [41, 42]. Importantly, several of the identified biomarkers represent drug targets, some of which are already in clinical use or tested in clinical studies including androgen receptor (AR) targeting drugs, EZH2 inhibitors, matrix metallopeptidases (MMP), transthyretin (TTR) or the RET proto oncogene.

For prim/norm samples, we identified two biomarkers associated with folate cleavage (*FOLH1*) and transport (*SLC19A1*), which might indicate an important dependency of PCa on folate metabolism. In fact, the *FOLH1* gene encodes for the prostate specific membrane antigen PSMA, which is an established tracer for positron emission tomography/computed tomography (^68^Ga PSMA PET/CT) and for radionuclide therapy (^177^Lu PSMA) [4]. In the met/prim we additionally identified *ADAM12*, a member of the A disintegrin and metalloproteases (*ADAM*) protein family, which is frequently up-regulated in solid cancers and involved in cell migration and invasion [43].

Cell surface proteins such as receptors or transporters represent suitable targets for specific diagnostic and therapeutic applications such as molecular imaging or targeted therapies. Therefore, we utilized the Cell Surface Protein Atlas tool (CSPA) [27] in order to identify up-regulated cell surface proteins (CSPs) in the pairwise comparisons described above. The CSP analysis resulted in a total of 246 proteins, of which 201 were significantly up-regulated in met/prim tumors and 45 were significantly up-regulated in prim/norm samples (Additional file 08). From those, 22 target genes were found to overlap in both comparisons, suggesting an important role of these factors for tumor progression and potentially prognosis. Metastatic tumors showed top up-regulated CSPs including several cell adhesion proteins, proteins regulating cell-cell and cell-matrix interactions or migration, as well as chemokine receptors (Figure 4A). The up-regulation of matrix associated factors including *OLFML2A/B*, *ADAM12*, *NTN3, LRRC15/32, CDH17, PCDH19, VTN, ICAM5* highlight the relevance of the tumor microenvironment for progression to metastatic disease. Interestingly, several of the identified genes are related to neuronal development or function including *NTSR1, NTN3, SERPINE2, SERPINI1*.

**Figure 4.**
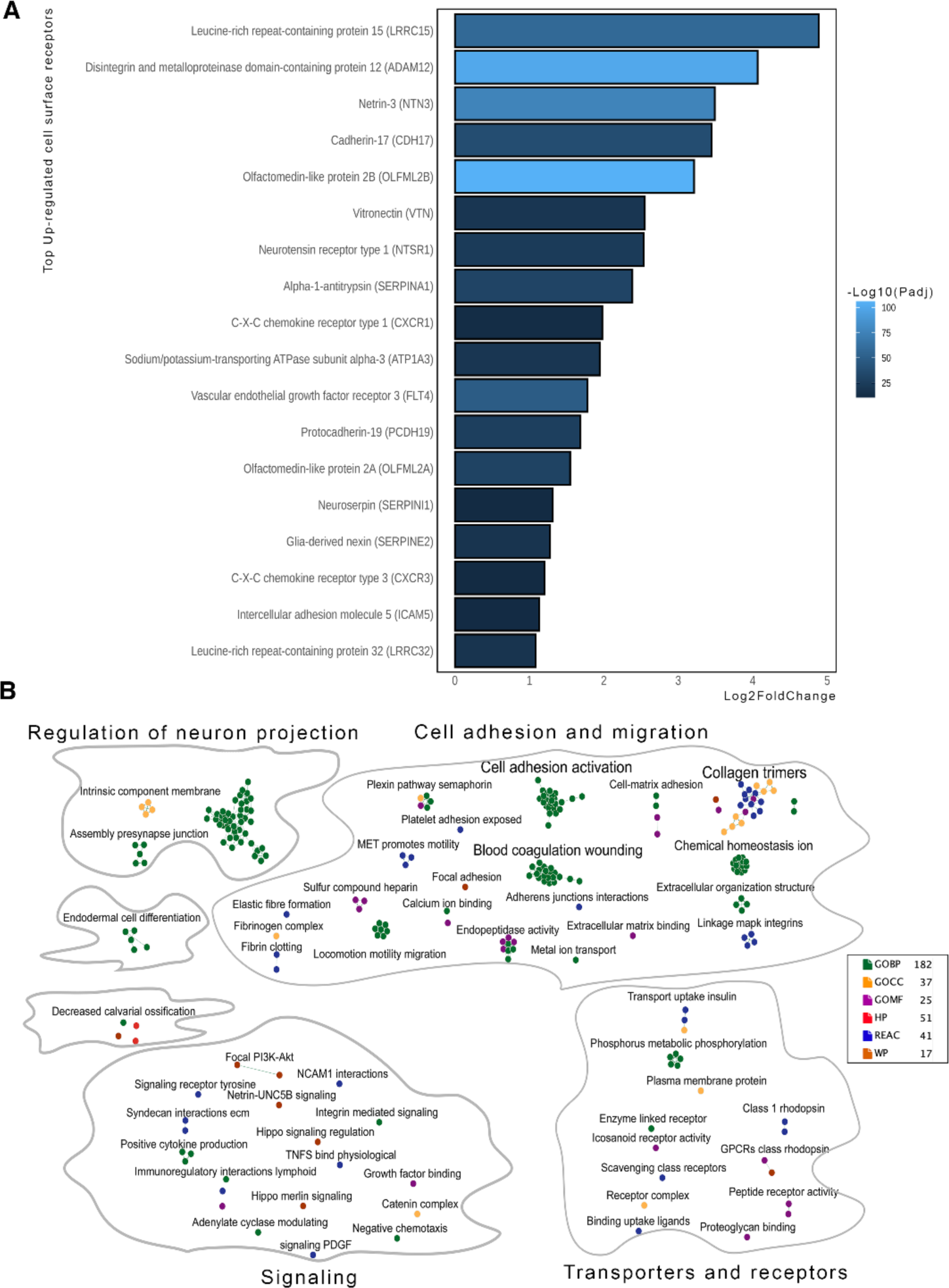
Integrative pathway enrichment analysis of cell surface protein receptors. **(A)** Cell surface proteins identified among the significantly up-regulated DEGs in met/prim samples, in which Padj are represented by colors and log2fold changes are represented by bar size. **(B)** Enrichment map of Gene ontology (GO) terms (GOBP, GO biological process; GOMF, GO molecular function; GOCC, Cellular Component), Human phenotype ontology (HP), WikiPathways (WP) and Reactome (REAC) pathways identified from up-regulated cell surface proteins in met/prim. Number of enriched terms in each database is shown on the enrichment map labels.

To gain further insight into the functional aspects of up-regulated CSPs in metastatic tumors, the identified genes were used for interactive pathway analysis [29]. The defined active pathways were used as input for the enrichment map analysis [31] of Cytoscape [32] for network visualization of similar pathways. Auto-annotated enrichment maps of up-regulated CSPs highlighted their potential functional implication for PCa. The pathways were grouped into four major classes including cell adhesion and migration, signaling, transporters and receptors and regulation of neuron projection (Figure 4B). Together these data suggest that PCa metastasis is highly dependent on proteins that are expressed at the surface of tumor cells or factors secreted into the extracellular space, which might represent efficient diagnostic and therapeutic targets for aggressive PCa.

### Identifying the most relevant DEGs using explainable machine learning yields multiple cancer-related biomarkers

In order to identify the most relevant DEGs to aid in biomarker identification, we trained explainable machine learning binary classifiers to distinguish between pairs of sample types, as defined by our experimental subgroups. Input DEGs were up-regulated (Padj < 0.05, LFC > 1) and uniquely mapped to ENTREZID/SYMBOL, were either protein-coding or non-coding RNA, and had overlapping expression values between the compared conditions. We used shapely additive explanations (SHAP) values [35], averaged over 10,000 bootstrap training iterations of random forests, to rank our up-regulated DEGs. SHAP values, which are generated for each sample and gene individually, indicate the impact that each gene has on the final model output, with higher absolute values demonstrating an increased influence. We achieved a mean test balanced accuracy among all bootstrap iterations of 0.95 (±0.05) for met/prim tumors and 0.86 (±0.07) for prim/norm samples.

Among the top ranked genes, we found relevant known cancer and PCa biomarkers for both examined comparisons, validating the suitability of our approach (Figure 5, A and B). For the prim/norm comparison, a total of 127 up-regulated DEGs above a SHAP threshold of 0.001 were determined (Supplementary Table S1, Additional file 09). Among the top genes, we identified known PCa genes such as *EPHA10, DLX2, HPN, HOXC4, HOXC6* and *AMACR*. *EPHA10* encodes for an ephrin receptor, whose overexpression in PCa might have potential as a target for therapy [44]. *DLX2* is a novel marker of increased metastasis risk [45]. *HPN* is a type II transmembrane serine protease and a marker to distinguish normal tissue from PCa lesion validated by immunostaining [46]. *HOXC4* and *HOXC6* are homeobox transcription factors commonly detected in PCa that co-localize with *HOXB13*, *FOXA1* and *AR*, three other transcription factors previously shown to contribute to the development of PCa [47]. Finally, the racemase *AMACR* is a known good predictor of clinically significant PCa [48]. For met/prim tumors, 117 up-regulated DEGs above a SHAP threshold of 0.001 were identified (Supplementary Table S2, Additional file 09). Genes related to other cancers such as *NPFF*, a neuropeptide and key biomarker for retinoblastoma [49], and *PMM2*, an enzyme involved in glycosylation and prognostic marker for colon cancer [50], were among the top hits. Further, cell surface receptors including *FGFRL1*, *LRFN1*, *NUP210*, *SMPDL3B* and *TMEM132A* for prim/norm and *ADAM12*, *MFAP3* and *YBX1* for met/prim were confirmed using the ML approach (Supplementary Table S4, Additional file 09).

**Figure 5.**
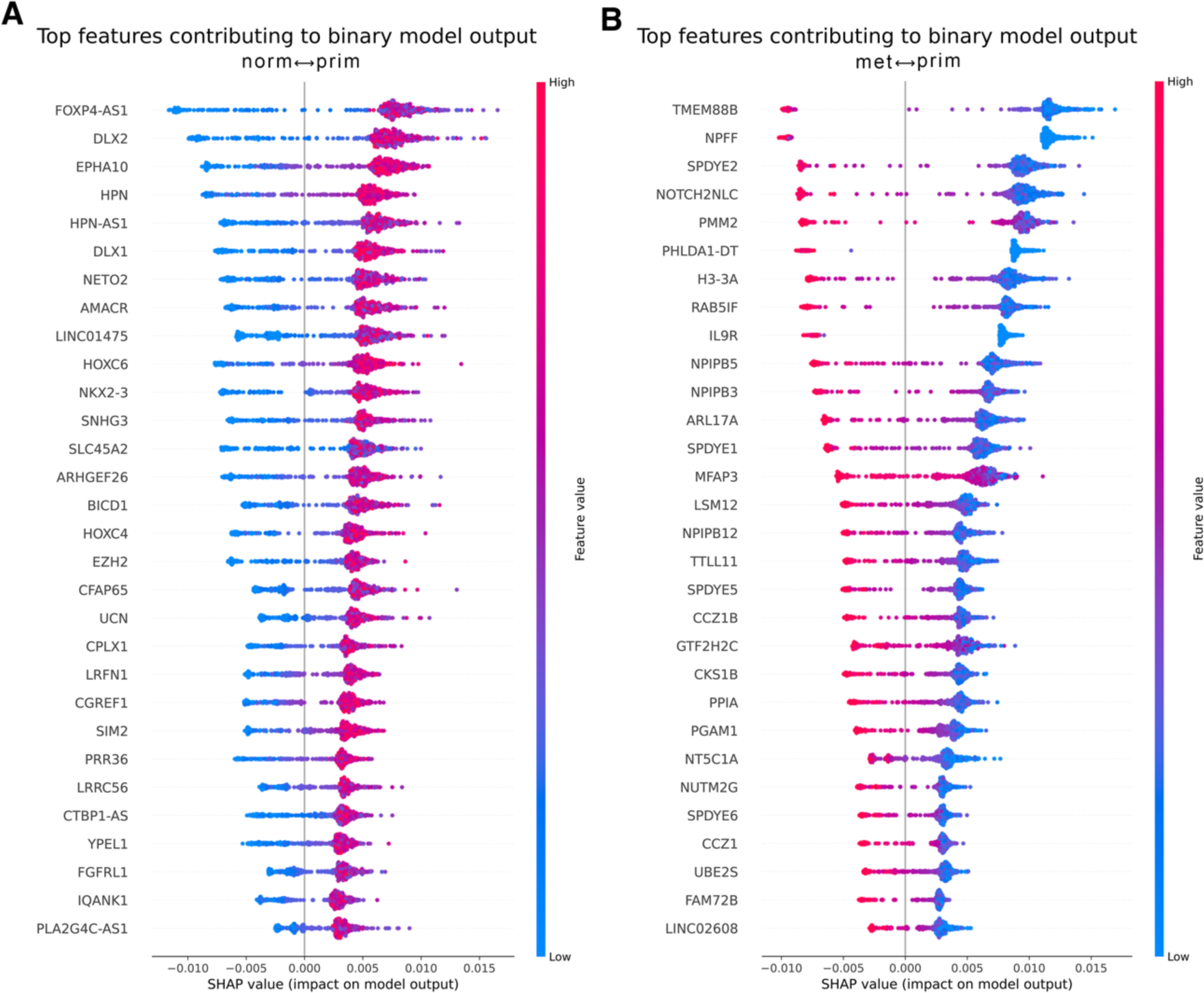
Beeswarm plots of explainable machine learning binary classifiers. Top differentially expressed genes (model features) influencing the output of explainable machine learning binary classifiers, as ranked by SHAP value, are shown for prim/norm **(A)** and met/prim **(B)**. Each dot represents a predicted sample color-coded by VST gene expression values (from blue, low expression to red, high expression). The further out a dot is from the separating vertical line, the higher the absolute SHAP value, as indicated in the X axis, and hence the higher the impact on the model output. Negative and positive SHAP values steer the model towards a classification output of 0 or 1, respectively.

Notably, *EZH2* and *TROAP* were identified in both group-wise comparisons, highlighting their potential as prognostic biomarkers. (Supplementary Table S3, Additional file 09). Both markers were recently also identified by employing ML tools on 30 previously published gene expression data sets [51].

Thus, our ML approach provided a significant number of candidate biomarker genes, some of which were already reported for their prognostic potential.

### Clustering of DEGs based on weighted gene co-expression networks serves to identify groups of similar biological processes related to PCa

Next, we analyzed co-expression relationships between up-regulated DEGs to get further biological insights, by grouping genes that act together in order to perform similar biological functions. Input DEGs were up-regulated (Padj < 0.05, LFC > 1) and uniquely mapped to ENTREZID/SYMBOL, were either protein-coding or non-coding RNA, and had both overlapping and non-overlapping expression values between the compared conditions.

For prim/norm, we identified a total of six modules of varying sizes as well as hub genes in each module according to their node degree (Figure 6A-B, Supplementary Tables S1-S6, Additional file 10). We were able to identify genes related to PCa among the module hub genes, such as *TPX2* in M3, which enables importin-alpha family protein and protein kinase binding activity and is highly expressed in high-grade PCa as well as being significantly related to poor prognosis [52]. Additionally, many of the most relevant genes in M3 were previously identified by our ML analyses and showed high SHAP values (Figure 5B), such as *HOXC6*, *AMACR* and *DLX2*, highlighting the relative biological significance of this module. Reactome functional enrichment analysis revealed enrichment of cell cycle and mitosis as the most significant pathways within the M3 module (Figure 6C, Additional file 11).

**Figure 6.**
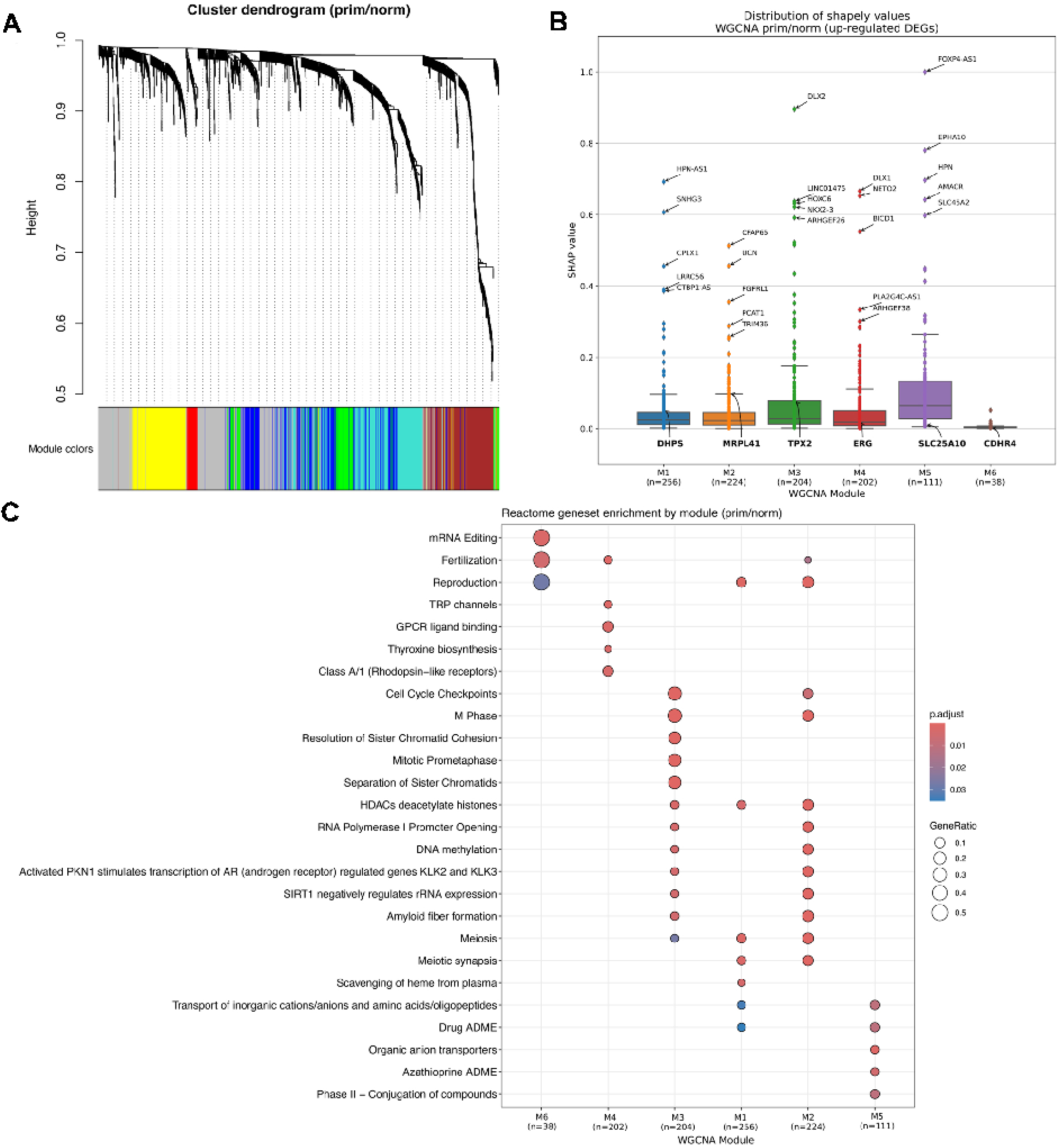
Weighted gene co-expression network analysis comparing primary versus normal samples. **(A)** Hierarchical cluster tree showing six modules of co-expressed genes. Each of the DEGs is represented by a leaf in the tree, and each of the nine modules by a major tree branch. The lower panel shows modules in designated colors, such as ‘red, ‘yellow, ‘brown’ and others. Note that module ‘grey’ is for unassigned genes. **(B)** Distribution of gene SHAP values for each WGCNA module. Top 5 genes with the highest SHAP values, as well as the module hub genes are highlighted. **(C)** Reactome functional enrichment results for each module, in which Padj are represented by colors and GeneRatio changes are represented by dot size.

For met/prim we identified a total of nine modules of varying sizes as well as hub genes in each module defined by node degree (Figure 7A-B, Supplementary Tables S1-S9, Additional file 12). Notably, *TPX2* was again detected as hub gene (M4), sharing 112 DEGs (26% of all genes present) with M3 in norm/prim (Supplementary Table S10, Additional file 12). Accordingly, Reactome analysis [25] revealed that M4 in met/prim contained all 23 pathways identified for M3 in prim/norm, and additional pathways associated with mitotic and cell cycle processes, as well as and pathways related to *TP53*. Moreover, in M4 the gene ratio in these shared pathways (i.e. the fraction of DEGs found in the gene set) was considerably bigger, which implied an increased activity of these pathways in metastatic tumors (Additional file 13). Together, these data highlight the potential of co-expression analysis for the identification of relevant modules that reflect major biological pathways promoting malignant transformation, tumor progression and metastasis. As a central theme, these analyses manifested the importance of transcriptional programs regulating mitosis and cell cycle control for PCa development and progression.

**Figure 7.**
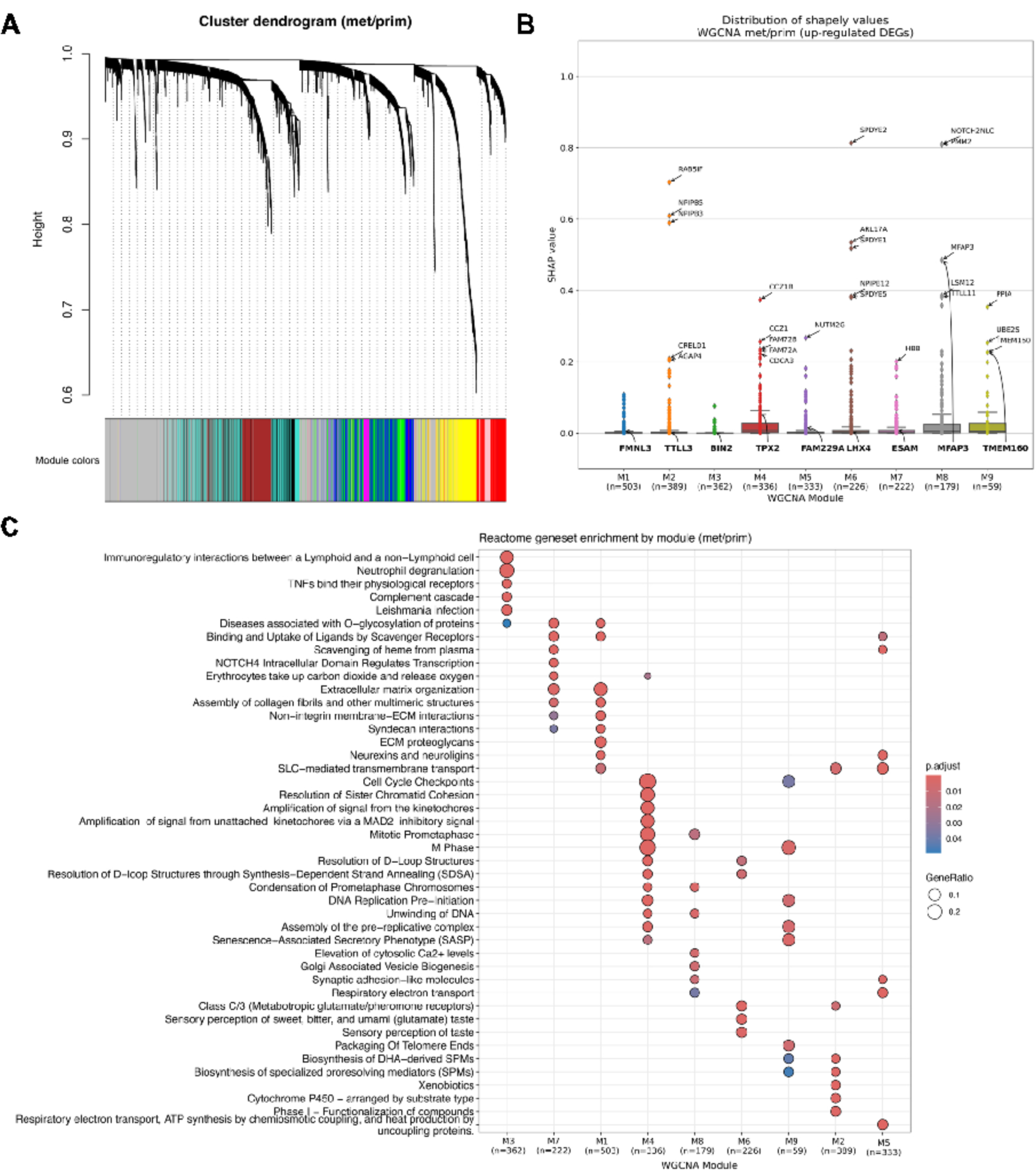
Weighted gene co-expression network analysis comparing metastatic versus primary tumors. **(A)** Hierarchical cluster tree showing nine modules of co-expressed genes. Each of the DEGs is represented by a leaf in the tree, and each of the nine modules by a major tree branch. The lower panel shows modules in designated colors, such as green, yellow, ‘turquoise’ and others. Note that module ‘grey’ is for unassigned genes. (B) Distribution of gene SHAP values for each WGCNA module. Top 5 genes with the highest SHAP values, as well as the module hub genes are highlighted. **(C)** Reactome functional enrichment results for each module, in which Padj are represented by colors and GeneRatio changes are represented by dot size.

### Integrative analysis for the mining of PCa candidate biomarkers

To obtain a conclusive, robust and more specific list of potential PCa biomarkers, we combined the outputs of the WGCNA and ML approaches. We focused on WGCNA modules M3 (prim/norm) and M4 (met/prim) given their overlap and previously discussed biological significance. First, we analyzed the overlap between the two modules and the ML results using a SHAP threshold of 0.001, which resulted in *EZH2* and *TROAP*, again highlighting the central role of these two proteins for PCa development and progression. Both genes, which were significantly up-regulated in the pairwise comparisons, showed a gradual increase of expression from normal to primary and metastatic samples (Figure 8A; Supplementary Table S1 Additional file 14).

**Figure 8.**
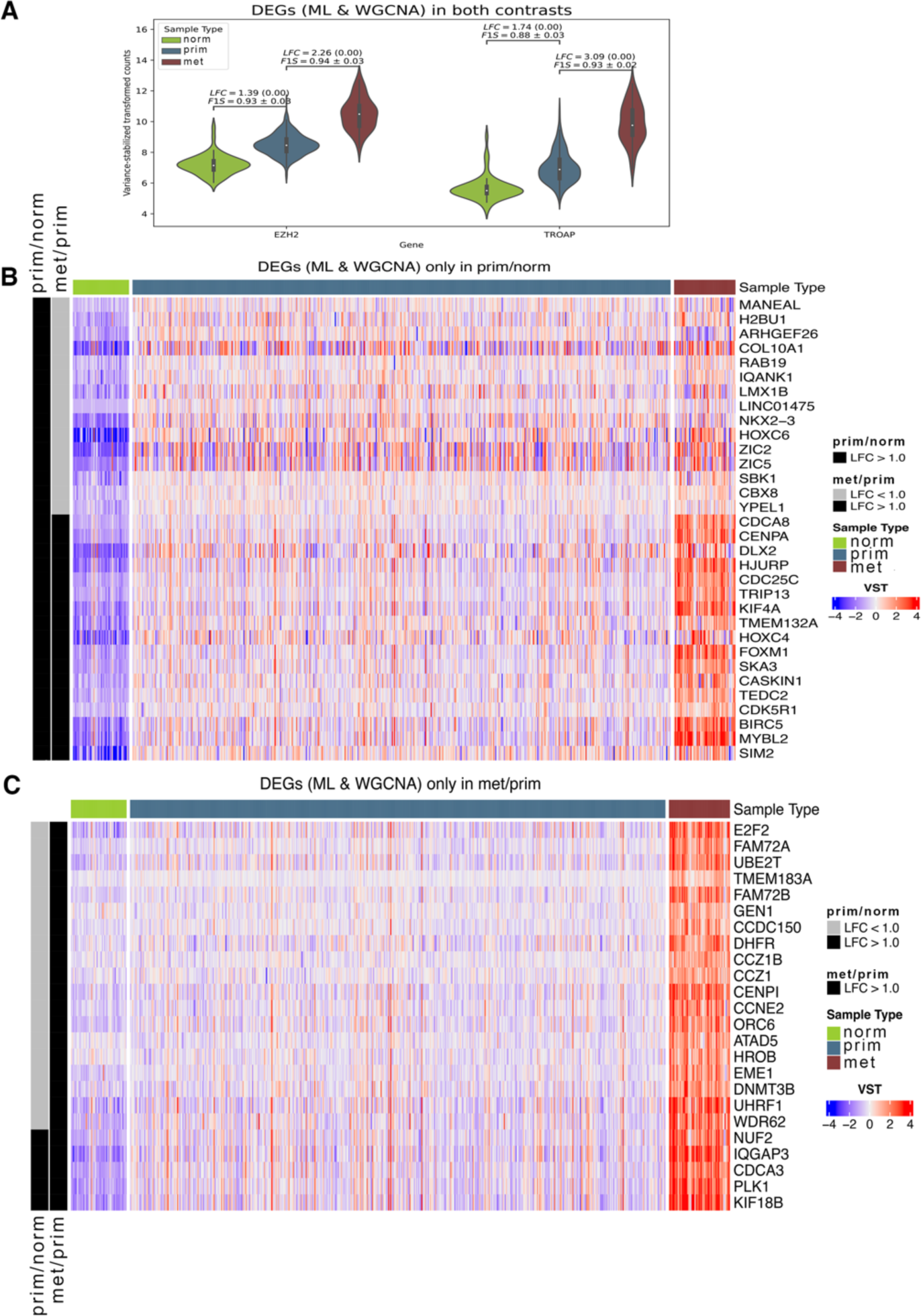
Candidate biomarker gene expression. **(A)** Violin plots showing intersecting DEGs (SHAP > 0.001) between prim/norm and met/prim filtered gene sets (WGCNA M3 and M4, respectively). LFC, Padj, and F1-score (F1S) metrics between comparisons are also shown. **(B)** Heatmap of DEGs (SHAP > 0.001) unique to the prim/norm filtered gene set (WGCNA M3), showing each DEG variance-stabilized transformed (VST) expression among the samples of each sample type. **(C)** Heatmap of DEGs (SHAP > 0.001) unique to the met/prim filtered set (WGCNA M4). In (B) and (C), for each contrast, samples are annotated on the left side of the heatmap as being biologically significant (LFC > 1) in black, or not, in gray.

To identify markers that were specific for primary PCa, we explored the SHAP-filtered DEGs in M3 (prim/norm) that were not present in module M4 (met/prim) (Supplementary Table S2, Additional file 14). We identified a total of 32 genes, 15 of which were not significantly up-regulated in met/prim (Figure 8B). Among these genes, we again found *HOXC6* as well as several other genes with known connections to PCa. Moreover, intersection of SHAP-filtered DEGs that were only present in M4 (met/prim) (Supplementary Table S3, Additional file 14), resulted in the identification of 24 genes, 19 of which were not significantly up-regulated in prim/norm (Figure 8C). We hypothesize that these genes, which were significantly up-regulated in metastatic tumors, are promising prognostic biomarker candidates.

Next, we performed *in silico* validation of the three inferred sets of biomarkers (*EZH2*,*TROAP* as progression markers, 32 candidate genes elevated in prim/norm, 24 candidate genes elevated in met/prim) on the Prostate Cancer Transcriptome Atlas (PCTA) dataset [37] after removing samples, which were already included in the initial analysis (i.e. those from TCGA-PRAD and SU2C-PCF) (Supplementary Tables S1-S3 Additional file 15, Additional file 16). We examined the differences in expression of the candidate biomarkers between both datasets and found a considerable rate of agreement. More concretely, we found that for both datasets, 74% of the biomarkers were similarly significantly up-regulated in prim/norm, whereas 79% were similarly significantly up-regulated in met/prim (Supplementary Tables S4-S5, Additional file 15).

Lastly, to determine the clinical relevance of these biomarkers, we performed recursive partitioning-based survival analysis [38] of the 58 individual biomarkers using the web-based camcAPP tool [40]. Using the MSKCC dataset [39], which includes primary and metastatic PCa samples, we found that 22 of the markers (addressed above) were significantly associated with shorter time to biochemical recurrence and thus worse prognosis of the patients (Additional file 17).

Thus, by integrating ML and WGCNA we were able to identify and validate different sets of candidate biomarkers, which are promising candidates for further evaluation in prospective clinical cohorts.

### Validation of *TPX2* as a potential target in PCa

As *TPX2* was a potential target in our differentially expressed analysis and was detected as hub gene in primary and metastatic PCa, we examined the protein expression of TPX2 on a tissue microarray (TMA) containing 51 primary PCa with adjacent normal tissues and 35 matched lymph node metastases (for TMA details see Methods section). Notably, while normal adjacent prostate epithelia were negative for TPX2 expression, a gradual significant increased expression was detected in primary PCa and lymph node metastases (Figure 9A). Together, these data confirm the upregulation of TPX2 during PCa progression on protein level (Figure 9B). Survival curves for high and low expression levels of TPX2 is significantly associated with shorter time to biochemical recurrence (Figure 9C), implying its biological role for tumor progression and its potential as a biomarker and therapeutic target for advanced PCa.

**Figure 9.**
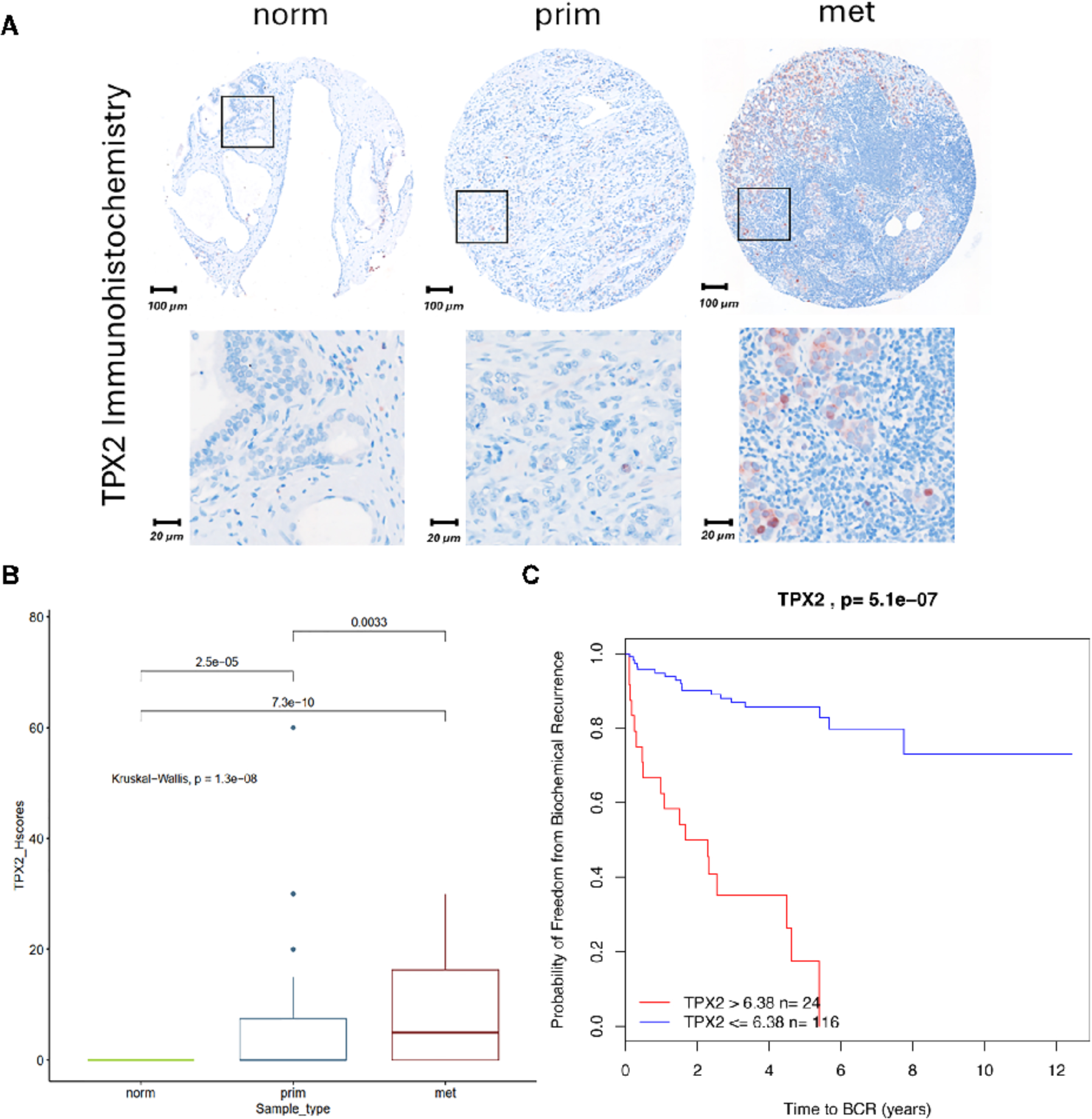
Validation of TPX2 protein expression. **(A)** Representative microscopic images of IHC stainings of normal (norm), primary PCa (prim) and lymph node metastatic (met) PCa samples. Brown nuclear staining indicates TPX2 expression. Sections were counterstained with hematoxilin (blue colour). **(B)** Protein expression levels indicated by H-scores, calculated from tissue microarrays containing 154 norm, 194 prim and 54 met. tissue core biopsies. **(C)** Kaplan Meyer survival curves for high and low expression levels of TPX2 is significantly associated with shorter time to biochemical recurrence.

## 4. Discussion

Tumor classification based on gene expression data revealed distinct PCa tumor subtypes and uncovered expression patterns that were associated with clinical outcomes for localized and metastatic disease [53, 54]. Thus, DEGs can serve as biomarkers for tumor diagnosis, prognosis, response prediction and/or as direct targets for drug development.

Our method extends transcriptomic analyses by using ML and WGCNA together with different pathway analysis tools to identify and characterize DEGs. This comprehensive approach allows for robust group prediction (norm/prim/met) and provides a deeper understanding of the functional roles of target genes implicated in PCa progression. By integrating datasets from the TCGA and dbGAP databases, we focused the analyses on genes that were up-regulated in primary localized PCa compared to benign tissue, or overexpressed in mCRPC compared to primary PCa. This unbiased setting provided the opportunity to include disease stages from localized to lethal PCa, thereby highlighting potential general biomarkers for PCa diagnosis (gene up-regulated in prim) or disease progression (genes up-regulated in met) irrespective of their heterogeneity or molecular subgroup.

Based on recent studies, cell surface proteins, which connect intracellular and extracellular signaling networks and largely determine a cell’s capacity to communicate and interact with its environment, are highly suitable biomarker candidates and targets for pharmacological intervention. However, information on the cell surface proteins of PCa is largely missing [27]. In our analysis, we identified a plethora of cell-surface proteins encompassing receptors, transporters, membrane-related proteins, cell-adhesion proteins, and extracellular enzymes as a rich source for biomarker development and as potential therapeutic targets. Notably, several top deregulated genes in metastatic tumors were associated with cell-cell or cell-matrix interactions, underscoring their relevance in metastatic processes.

Interestingly, our analysis revealed deregulation of genes associated with neuronal processes among the top DEGs. For instance, *NTN3* (a gene part of the nexin family of proteins) and *NTSR1* (neurotensin receptor 1) have been linked to the progression of PCa. *NTN3* regulation by the AR-EZH2 axis in PCa has been documented [55], while NTSRs and its ligand neurotensin (NTS) critically affect the progression of PCa. Immunohistochemical analysis have indicated high or moderate expression of NTSR1 in 91,8% of PCa tissue including PSMA negative samples [56] and in 9,1% in primary PCa and 33% of lymph node metastases [57], in two recent studies. In addition, molecular imaging tracers are available and might represent suitable tracers for PET imaging and radionuclide therapy, especially for PSMA negative tumors or tumors with neuroendocrine differentiation [58]. Together, these data imply important functions for several neurodevelopmental genes and pathways for PCa progression also in non-neuroendocrine differentiated tumors.

Our integrative approach, which combined ML and WGCNA, revealed a consistent trajectory towards disease progression from normal prostate through primary to metastatic PCa involving a gradual up-regulation *EZH2* and *TROAP.* This underscores their pivotal roles in PCa progression. *TROAP* encodes for the trophinin associated protein that has been found to regulate PCa progression via the WNT3/survivin signaling pathways due to its essential role in cell proliferation [59]. Interestingly, *TROAP* was shown to be downstream of *EZH2* and is important for PCa progression via the TWIST/c-MYC pathway [60]. The role of the polycomb repressive complex PRC2 protein *EZH2* has been well established for PCa progression to mCRPC or lineage plasticity of neuroendocrine PCa [61, 62]. EZH2 also acts in a PRC2 independent fashion as an *AR* co-activator or by transcriptional activation of *AR* [63]. *EZH2* is currently assessed as a therapeutic target for mCRPC in clinical studies, using the EZH2 inhibitor tazemetostat in combination with androgen signaling inhibitors (NCT04179864) or the PARPi talazoparib (NCT04846478).

In addition, the integration of ML and WGCNA yielded a selection of candidate biomarkers for localized PCa, including *SBK1*, *ARHGEF26*, *DLX2* and *NKX2*-3, or *HROB*, *GEN1* and *E2F2* for mCRPC, whose clear role for PCa has only been partially elucidated. This, might warrant further functional studies, especially for genes up-regulated in metastatic tumors including *HROB* or *GEN1*, which are implicated in DNA damage repair pathways or *SBK1*, *DLX2* and *E2F2* which are associated with worse prognosis of patients.

In addition to *EZH2* and *TROAP*, through WGCNA analysis, we identified key co-expression modules and hub genes critical in several cancer-related pathways. Among several relevant genes, *TPX2* was identified as a hub gene for both pairwise comparisons, highlighting its relevance for PCa development and progression. Our study identified *TPX2* as a crucial player in PCa development and progression [64], suggesting its potential as a prognostic marker and therapeutic target. The roles of TPX2 in DNA repair mechanisms and its interaction with PARP1 further underscore its importance in PCa biology. TPX2 is an activator of AURKA signaling and is important for microtubule nucleation and stability during mitosis [65]. AURKA is regulated by AR and represents a therapeutic target for PCa [66]. Furthermore, TPX2 is involved in homology-directed repair of DNA double strand breaks during replication stress and is a direct interaction partner of PARP1 [67]. This might also have important implications for PARP inhibitor therapy for advanced PCa. Similar to our analysis, a recent WGCNA analysis of the TCGA PRAD dataset identified *TPX2* as a hub gene and confirmed its prognostic value [68].

Future studies aimed at elucidating the mechanistic roles of TPX2 in PCa pathogenesis and evaluating its clinical utility are warranted to harness its full potential in improving patient outcomes and guiding personalized treatment strategies for prostate cancer.

## Conclusion

This work emphasizes the gradual transcriptional adaptations from primary to metastatic PCa and highlights several relevant disease pathways, drug targets and candidate biomarker genes. Future work that integrates multi omics data such as mutation data, epigenomics and RNA-Seq data in large cohorts will enable further insights into the identified biomarkers. In parallel, complementary experimental approaches to study the functional relevance of the identified candidate markers will provide further information on potential targets for therapeutic intervention for PCa and to translate promising candidates into clinical routine.

## List of abbreviations

PCa: Prostate cancer
mCRPC: Metastatic castration-resistant PCa
BCR: Biochemical recurrence
PSA: Prostate specific antigen
DDR: DNA damage repair
MMR: DNA mismatch repair genes
PSMA: Prostate-specific membrane antigen
TCGA: The Cancer Genome Atlas
PRAD: Prostate adenocarcinoma dataset
dbGap: Database of Genotypes and Phenotypes
prim: Primary PCa
norm: Normal adjacent tissues
met: Metastatic PCa
DE: Differential gene expression analysis
Padj: Adjusted P-value
LFC: Log2 fold change
VST: Variance stabilizing transformation
DEGs: Differentially expressed genes
GO: Gene ontology
CSP: Cell surface protein
CSPA: Cell surface protein atlas
HP: Human phenotype ontology
WP: WikiPathways
ML: Machine learning
SHAP: SHapley Additive exPlanations
WGCNA: Weighted correlation network analysis
PCTA: Prostate Cancer Transcriptome Atlas
IPA: Ingenuity Pathway Analysis
PPI: Protein-protein interaction
FFPE: Formalin-fixed paraffin-embedded
TMA: Tissue microarrays
IHC: Immunohistochemical

## Declarations

### Ethics approval and consent to participate

Not applicable

### Consent for publication

Not applicable

### Availability of data

Data used in this study were extracted from the TCGA Prostate Adenocarcinoma project (TCGA-PRAD, https://portal.gdc.cancer.gov/repository) and the SU2C-PCF dataset via dbGAP, with accession number phs000915.v2.p2 (https://www.ncbi.nlm.nih.gov/projects/gap/cgi-bin/study.cgi?study_id=phs000915.v2.p2).

### Authors’ contributions

GE and RST conceived and designed the study and drafted the manuscript. CUPM and RST performed all computational analyses. GE, RST and CUPM were major contributors in writing the manuscript. AM collected the patient data from public databases and commented on the manuscript. KM, LT, JK and GW contributed in background and literature search and editing the manuscript. All authors read and approved the final manuscript.

## Acknowledgements

The results shown in our research are in whole based upon data generated by the TCGA Research Network (https://www.cancer.gov/tcga) and and the database of Genotypes and Phenotypes (dbGAP) (https://www.ncbi.nlm.nih.gov/gap/). The authors would like to thank for the availability of TCGA data and also getting authorized access to dbGAP database in this study. This work was supported by grants provided by the FWF (National Science Foundation in Austria, grant no. P 32771) and DOC Fellowship of the Austrian Academy of Sciences (25276).

## Authors information

**1. Ludwig Boltzmann Institute Applied Diagnostics, Vienna, Austria**

Raheleh Sheibani-Tezerji, Carlos Uziel Pérez Malla, Anna Malzer, Katarina Misura, Loan Tran & Gerda Egger

**2. Department of Pathology, Medical University of Vienna, Vienna, Austria**

Raheleh Sheibani-Tezerji, Jessica Kalla, Anna Malzer, Katarina Misura, Loan Tran, Gabriel Wasinger, Astrid Haase & Gerda Egger

***3.* Comprehensive Cancer Center, Medical University of Vienna, Vienna, Austria**

Gerda Egger

## Description of Additional Files

**Additional File 1. Principal component analysis (PCA) for all samples from TCGA and dbGAP databases.** PCA graph of RNA expression levels for metastatic samples with unique patient Ids (Metastatic_AA: n=67, Metastatic_BB: n=57, Endocrine: n=14), primary tumors (n=500) and normal (n=52) samples from patients with PCa and mCRPC obtained from dbGAP and TCGA databases.

**Additional File 2. Annotation summary.** Annotation data and clinical information of all 609 Metastatic, primary and normal samples.

**Additional File 3. Principal component analysis (PCA) of RNA expression levels.** PCA graphs including PC1-4 show sample variance. Individual samples (circles) are color-coded by sample type.

**Additional File 4. Differentially expressed genes. Supplementary Table S1:** Differentially expressed genes (Padj < 0.05, abs (LFC) ≥ 1) for prim/norm (only unique ENTREZID and SYMBOL). **Supplementary Table S2:** Differentially expressed genes (Padj < 0.05, abs (LFC) > 1) for met/prim (only unique ENTREZID and SYMBOL). **Supplementary Table S3:** Shared up-regulated DEGs (Padj < 0.05, abs (LFC) > 1) between experimental subgroups. **Supplementary Table S4:** Shared down-regulated DEGs (Padj < 0.05, abs (LFC) > 1) between experimental subgroups

**Additional File 5. Gene ontology (GO) enrichment analysis in prim/norm.**

**Supplementary Table S1:** GO ontology enrichment analysis of significantly differentially expressed genes in prim/norm. **Supplementary Table S2:** GO ontology enrichment analysis of significantly differentially expressed genes in met/prim

**Additional File 6. Ingenuity Pathway Analysis (IPA) in prim/norm.**

**Supplementary Table S1:** Significant IPA pathways in up-regulated DEGs from prim/norm. **Supplementary Table S2:** Significant IPA pathways in up-regulated DEGs from met/prim.

**Additional File 7. IPA biomarker analysis in prim/norm.**

**Supplementary Table S1:** Potential biomarkers and targets in up-regulated DEGs from prim/norm. **Supplementary Table S2:** Potential biomarkers and targets in up-regulated DEGs from met/prim. **Supplementary Table S3:** Shared up-regulated biomarkers between experimental subgroups.

**Additional File 8. Cell surface proteins analysis in prim/norm.**

**Supplementary Table S1:** Top cell surface proteins (based on significant differential gene expression in met/prim) in up-regulated DEGs from prim/norm. **Supplementary Table S2:** Top cell surface proteins (based on significant differential gene expression in met/prim) in up-regulated DEGs from met/prim. **Supplementary Table S3**: Shared up-regulated cell surface proteins between experimental subgroups.

**Additional File 9. Top prim/norm up-regulated genes filtered by SHAP values. Supplementary Table S1:** Up-regulated DEGs (Padj < 0.05, LFC > 1, prim/norm) with SHAP > 0.001 **Supplementary Table S2:** Up-regulated DEGs (Padj < 0.05, LFC > 1, met/prim) with SHAP > 0.001. **Supplementary Table S3:** Overlapping Up-regulated DEGs (Padj < 0.05, LFC > 1) with SHAP > 0.001 between prim/norm and met/prim. **Supplementary Table S4:** Overlapping Up-regulated DEGs (Padj < 0.05, LFC > 1) with SHAP > 0.001 between prim/norm, met/prim and validated ML signatures.

**Additional File 10. WGCNA analysis genes for prim/norm. Supplementary Table S1:** WGCNA M1 DEGs (Padj < 0.05, LFC > 1, prim/norm). **Supplementary Table S2:** WGCNA M2 DEGs (Padj < 0.05, LFC > 1, prim/norm). **Supplementary Table S3:** WGCNA M3 DEGs (Padj < 0.05, LFC > 1, prim/norm). **Supplementary Table S4:** WGCNA M4 DEGs (Padj < 0.05, LFC > 1, prim/norm). **Supplementary Table S5:** WGCNA M5 DEGs (Padj < 0.05, LFC > 1, prim/norm). **Supplementary Table S6:** WGCNA M6 DEGs (Padj < 0.05, LFC > 1, prim/norm)

**Additional File 11. Enriched Reactome pathways in WGCNA.**

**Supplementary Table S1:** WGCNA Reactome enriched pathways in all modules (Padj < 0.05, LFC > 1, prim/norm).

**Additional File 12. WGCNA analysis genes for met/prim.** Met/prim up-regulated differentially expressed genes (DEGs), filtered by statistical and practical significance (i.e. Padj < 0.05, LFC > 1), as identified by WGCNA and presented in modules 1 to 9 (**Supplementary Tables S1 to S9**). **Supplementary Tables S10:** Overlapping up-regulated differentially expressed genes (DEGs), filtered by statistical and practical significance (i.e. Padj < 0.05, LFC > 1), shared between WGCNA module 3 of prim/norm and WGCNA module 4 of met/prim. The second and third columns indicate in which module the DEGs were present.

**Additional File 13. Enriched Reactome pathways in WGCNA.**

**Supplementary Table S1:** WGCNA Reactome enriched pathways in all modules (Padj < 0.05, LFC > 1, met/prim).

**Additional File 14. Shared and unique biomarkers from WGCNA.**

**Supplementary Table S1:** Shared biomarkers (prim/norm WGCNA M3, met/prim WGCNA M4, SHAP > 0.001). **Supplementary Table S2:** Prim/norm only biomarkers (prim/norm WGCNA M3, SHAP > 0.001). **Supplementary Table S3:** Met/prim only biomarkers (met/prim WGCNA M4, SHAP > 0.001).

**Additional File 15. Shared biomarkers validated in PCTA.**

**Supplementary Table S1:** Up-regulated differentially expressed genes (DEGs), filtered by statistical and practical significance (i.e. Padj < 0.05, LFC > 1), with a SHAP value bigger than 0.001 and shared between WGCNA module 3 in prim/norm and WGCNA module 4 in met/prim. Differential expression and predictive metrics are computed on the PCTA dataset for validation. **Supplementary Table S2:** Up-regulated differentially expressed genes (DEGs), filtered by statistical and practical significance (i.e. Padj < 0.05, LFC > 1), with a SHAP value bigger than 0.001 and unique to WGCNA module 3 in prim/norm. Differential expression and predictive metrics are computed on the PCTA dataset for validation. **Supplementary Table S3:** Up-regulated differentially expressed genes (DEGs), filtered by statistical and practical significance (i.e. Padj < 0.05, LFC > 1), with a SHAP value bigger than 0.001 and unique to WGCNA module 4 in met/prim. Differential expression and predictive metrics are computed on the PCTA dataset for validation. **Supplementary Table S4:** Combined differential expression and predictive metrics computes on our data and the PCTA dataset. The first column indicates the biomarker set (out of three possible), whereas the second column indicates in which dataset the metrics were computed. **Supplementary Table S5:** Comparison summary between the differential expression metrics on our data and the PCTA dataset. Cells highlighted in green indicate biomarkers that are equally statistically and significantly expressed, whereas red cells indicate some difference in expression patterns between both datasets.

**Additional File 16. Validation of candidate biomarkers on Prostate Cancer Transcriptome Atlas (PCTA)**. Only up-regulated DEGs (Padj < 0.05, LFC > 1) were considered. **(A)** Violin plots show intersecting DEGs (SHAP > 0.001) between prim/norm and met/prim filtered gene sets (WGCNA M2 and M4, respectively). LFC, Padj, and F1-score (F1S) metrics between comparisons are also shown. **(B)** Heatmap of DEGs (SHAP > 0.001) unique to the prim/norm filtered gene set (WGCNA M2), showing each DEG variance-stabilized transformed (VST) expression among the samples of each sample type. **(C)** Heatmap of DEGs (SHAP > 0.001) unique to the met/prim filtered set (WGCNA M4). In B) and C), for each contrast, samples are annotated on the left side of the heatmap as being biologically significant (LFC > 1) in black, or not, in gray.

**Additional File 17. Kaplan Meyer survival curves for high and low expression levels of the 22 biomarkers were significantly associated with shorter time to biochemical recurrence. (A)** Survival curves of intersecting DEGs (SHAP > 0.001) between prim/norm and met/prim (WGCNA M2 and M4, respectively). **(B)** Survival curves of unique DEGs (SHAP > 0.001) to the prim/norm filtered gene set (WGCNA M2). **(C)** Survival curves of unique DEGs (SHAP > 0.001) to the met/prim filtered set (WGCNA M4).

